# Deep Bidirectional Recurrent Neural Networks as End-To-End Models for Smoking Status Extraction from Clinical Notes in Spanish

**DOI:** 10.1101/320846

**Authors:** Santiago Esteban, Manuel Rodríguez Tablado, Francisco E. Peper, Sergio A. Terrasa, Karin S. Kopitowski

## Abstract

**Introduction:** Although natural language processing (NLP) tools have been available in English for quite some time, this is not the case for many other languages, particularly for texts from specific contexts such as clinical texts. This poses a challenge for tasks such as classifying text in languages other than English. In the absence of basic NLP tools, the development of statistical models that include manually designed variables that capture the semantic information of the documents is a potential solution. However, this process is expensive and slow. Deep recurrent neural networks (RNNs) have been proposed as “end-to-end” models that learn both variables and parameters jointly, thus avoiding manual feature engineering and saving development time.

**Methods:** We compared the performance of two strategies for labeling clinical notes of an electronic medical record in Spanish according to the patient’s smoking status (current smoker, current non-smoker, text without information on tobacco): 1. A traditional approach using two classifiers (a multilayer perceptron (MLP) and a support vector machine (SVM)) together with a ‘bag-of-words’ text representation that involves intensive manual development of features and, 2. an ’end-to-end’ model which uses a Short-Long-Term Memory bidirectional deep RNN with GloVe word embeddings. The classifiers were trained in the training set (n = 11775 clinical texts) and were evaluated in the test set (n = 2943) by means of macro-averaged recall, precision and F1 score.

**Results:** The RNN scored high values of all three metrics in the test set (sensitivity [95% CI]: 0.965 [0.96, 0.97], PPV: 0.963 [0.96, 0.97], F1 score: 0.964 [0.96, 0.97]). It also showed to be slightly superior to the MLP (difference in recall: 0.009 [95% CI: -0.0007, 0.017], precision: 0.007 [95% CI: -0.0015, 0.019] and F1 score: 0.009 [95% CI: 0.0018, 0.016]); comparing the RNN with the SVM, the latter has a better performance in general (recall difference [95% CI]: -0.007 [-0.016, 0.0018], precision: -0.009 [-0.018, 0.00015] and score F1: -0.008 [-0.014, -0.0017]). In both cases only the confidence interval for the F1 score difference excludes zero. In turn, the RNN consumed 80% less overall development time.

**Conclusion:** In our work, the deep bidirectional RNN as end-to-end model, reached similar levels of performance in the classification of clinical texts in Spanish that models with a great manual engineering of variables, although in less than 20% of the development time. This makes them an important tool to streamline text processing in languages where the development of NLP tools has not progressed as much as in English. Areas such as research or public health management could clearly benefit from ’end-to-end’ models that facilitate the exploitation of already available data sources, such as electronic clinical records.

## Introduction

Globally, tobacco is one of the main risk factors for premature death^[1]^ being responsible for 11.5% of the worldwide annual deaths (6.5 million)^[2]^. It is also one of the five main risk factors of disability^[2]^. In 2015, the global prevalence of daily tobacco use was estimated at 25% (uncertainty interval (UI) 24.2-25.7) in men and 5.4% (UI 5.1-5.7) in women with a marked decrease since 1990^[2]^. However, despite these general trends, there is a high level of heterogeneity among countries and this decline was not as marked in countries with low and middle socio-demographic levels, especially in women^[2]^. This is evident in that of the 933 million daily smokers, 80% reside in such countries^[3]^. However, despite its importance, clinical and public health research on risk factors such as smoking is often restricted, among other things, by the scarcity of trained human resources and economic resources^[4, 5]^.

In this context, electronic health records (EHR) are proposed as data sources that facilitate clinical research, especially taking into account the increasing adoption rate in both developed and developing countries^[6]^.

Information regarding tobacco consumption is usually recorded in the EHRs in different formats, both as structured problems but also as free text, usually complementing each other^[7]^. However, there is a clear tendency to register a large part of the relevant information on smoking in the free text due to, among other reasons, the lack of flexibility of the more structured recording systems^[8]^. This is why many authors have explored methods based on natural language processing (NLP) for the extraction of information regarding the patients’ smoking status from clinical notes^[9–17]^.

The task of classifying texts by assigning them a predefined category or label (for example: smoker, non-smoker, ex-smoker) has traditionally been solved using techniques based on hard-coded rules and techniques based on statistical or supervised machine-learning^[18]^, which have progressively dominated NLP tasks in the last 30 years.

One of the most disseminated and also simplest approaches is the bag-of-words model^[19]^. In this model each gram (term or token) or n-gram (concatenation of n tokens) of the corpus (collection of documents) is treated as a feature and a value is assigned to each feature based on the weighted or unweighted counts of the terms in each document. Thus, each document is represented by a feature vector of these counts. In turn, all the feature vectors compose a document-term-matrix, where each observation (document) is a row and each feature, a column. Once this matrix is created, a classifier algorithm is iteratively trained to minimize a cost function in order to find the best separation plane between the categories and predict de document’s label. The performance of this method can be improved by developing variables that better capture the linguistic structure of the text. Many NLP tools have been designed to create such features like constituency^[20]^ and dependency parsing^[21]^, negation detection^[22, 23]^, part-of-speech (POS) tagging^[24]^ or named-entity recognition^[25]^ tools.

However, these tools are, for the most part, language-specific and, in many cases, context-specific. This means that, in order to apply them in a language or context different from the one they were developed in, it is necessary to retrain (re-learn the parameters of the model in a new dataset) or even redesign them completely. Clinical notes from EHRs are clearly different from usual texts: they contain specific jargon, incomplete or grammatically incorrect sentences and a large number of abbreviations and acronyms. This implies that, in order to develop the beforementioned NLP tools for specific languages and contexts, it is necessary to develop a training corpus, usually annotated by linguistics experts (“gold corpus”), which is an expensive and slow process that in many cases ends up being prohibitive in terms of resource expenditure. This aspect is rather important considering that most of the research and tools in NLP have been developed in English and most of the developing countries, where it would be essential to promote research on smoking, do not have English as their primary language and therefore, these tools are not available in an “off-the-shelf” fashion.

A potentially more accessible solution is the extension of the models described so far by developing simple linguistic models and manually engineering features specific to the classification task. This is usually done by rigorous exploration of the text and with specific knowledge of the classification problem itself. Examples of this would be the identification of key topic words, the development of specific tools to detect key term negations and modifiers. This, although feasible, requires the investment of many hours in the process of manual feature engineering and feature selection. This approach, nevertheless, suffers from the fact that the models are not usually exportable to other classification problems.

### End-to-End models for text classification

The models described so far are usually implemented in two phases, where the features and the parameters of the models are developed separately and by different actors: the features are manually engineered by humans, the parameters are estimated from the data by the classifier. The philosophy behind ‘End-to-End’ models seeks that both, features and parameters are learned jointly and by the same actor: the algorithm. In recent years ‘End-to-End’ models have been excelling at tasks such as image classification^[26]^, machine translation^[27]^, autonomous driving^[28]^ and speech recognition^[29]^. Most of these examples have used deep neural networks but the ‘End-to-End’ philosophy exceeds this type of algorithms. Basically, End-to-End models remove intermediate steps by freeing the human from the task of manually engineering features and can potentially help find patterns in the data that would be difficult for a human to discover. In the specific case of NLP, they can avoid having to develop many of the most widely used tools (parsers, POS taggers, etc.). This is particularly advantageous for languages or contexts for which prebuilt tools are not available or don’t perform very well.

### Bidirectional Long Short-Term Memory (LSTM) Recurrent Neural Networks (RNN) as End-to-End models for text classification

Recurrent neural networks are a type of artificial neural network architecture that allows to process series of data sequentially. Basically, the hidden state of the previous neuron (*h*_*t*−1_) is part of the input of the neuron *t*_0_ along with the input at *t* (*x*_*t*_) (fig. 1). Thus, *h*_*t*_ depends on *h*_*t*-1_ and *X*_*t*_ (*h*_*t*_ = *f*(*W*^*xh*^*x*_*t*_ + *W*^*hh*^*h*_*t*−1_ + *b*^*h*^)), where *f* is usually a non-linear activation function (sigmoid, tanh, ReLU) and *b* is the bias term. Unlike in feed-forward neural networks, the weights (*W*^*xh*^, *W*^*hh*^, *W*^*hy*^) are shared for each time step *t*. This sharing of weights is what gives RNN the ability to ‘remember’ the hidden states of previous neurons and allows modeling of data sequences by capturing sequential patterns. Nevertheless, in practice, standard RNN are not capable of handling long-term dependencies because of the vanishing and exploding gradients problem initially detailed by Bengio et al^[30]^. Modifications to the structure of basic RNNs (Long-Short Term Memory^[31]^ units; Gated Recurrent Units^[32]^) have allowed these algorithms to learn even more distant dependencies within the series without suffering from the problems detailed above. Being able to represent long term dependencies makes them natural candidates for the classification of texts, since they can represent key aspects of linguistic structures that resembles human text processing^[33]^. In addition to this, bdRNNs (bidirectional RNNs)^[34]^ allows for the processing of the texts such that *h*_*t*_ is also influenced by data further down the sequence. Also, stacking of such bidirectional RNNs in layers enables the network to learn hierarchical more abstract representations (fig. 2). Moreover, the representation of words by means of pretrained word embeddings^[35–37]^ further increases the representational capacity of these models.

**Figure 1.**
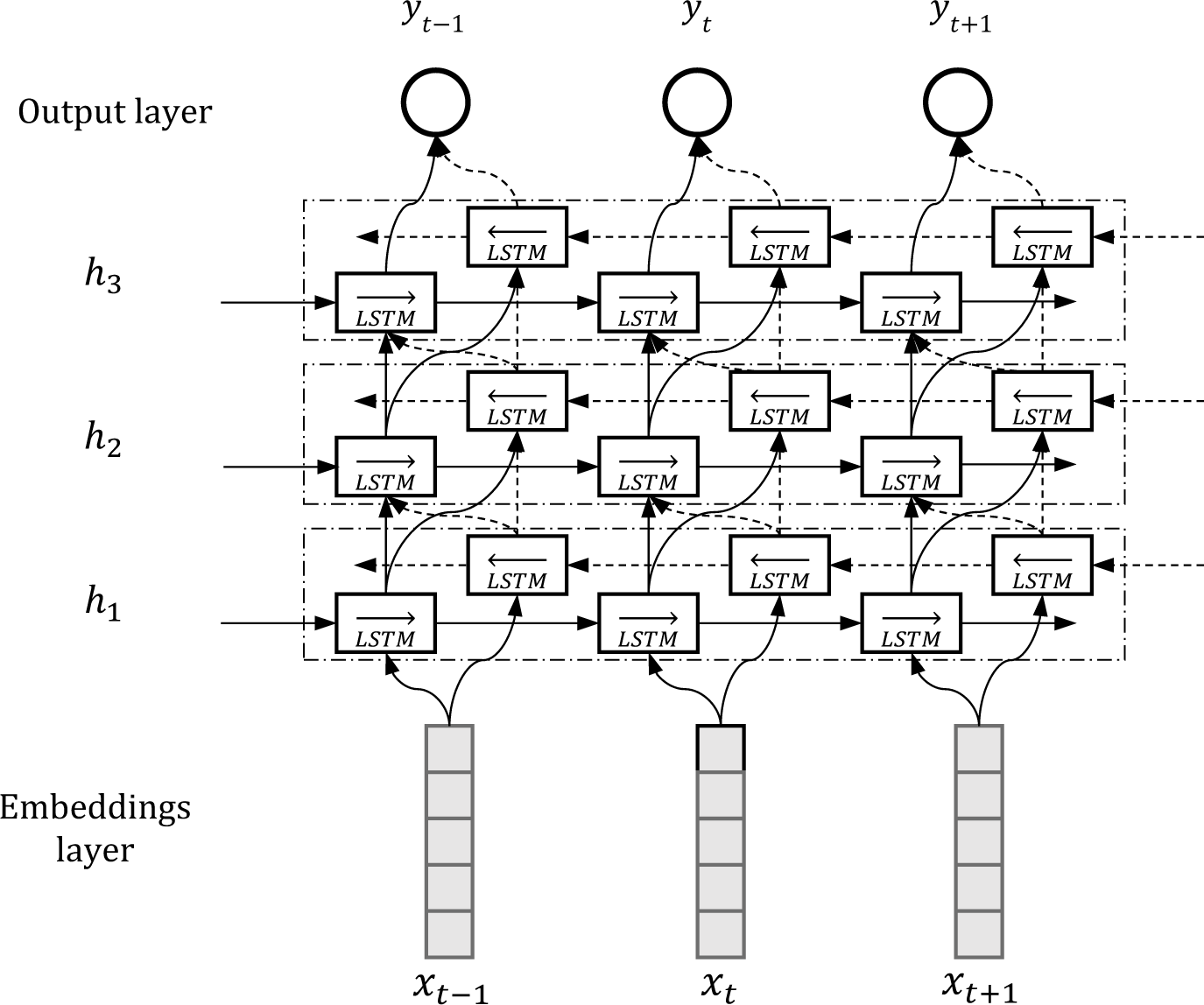
An unrolled recurrent neural network. For simplicity the biases are not depicted. *x*: input; *h*: hidden state; *t*: time step; *y*: output; *w*: weights.

**Figure 2.**
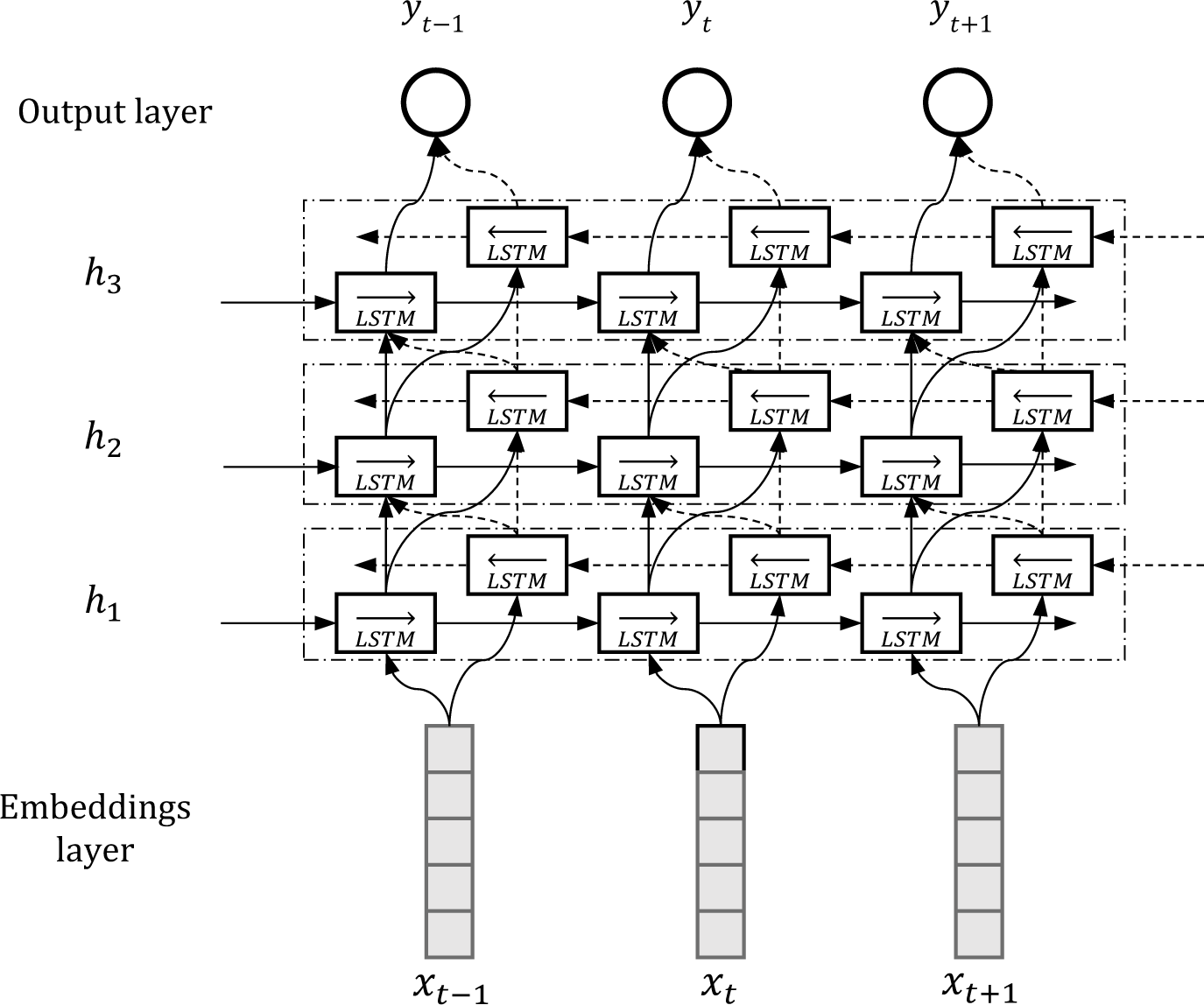
An unrolled bidirectional long-short-term-memory recurrent neural network. *LSTM*: long-short-term memory; *t*: time step; *h*: hidden layer; *x*: input; *y*: output.

## Background

In recent years, End-to-End models implemented by means of artificial neural networks (ANN) have become more relevant for NLP tasks in general ^[38–41]^ as well as in specific domains, like clinical narratives. In 2016, Jagannatha et al. ^[42]^ compared the performance of a bidirectional RNN vs conditional random fields (CRF)^[43]^ for detecting medical events in notes from an EHR. They found improvements in accuracy, recall and F1 score vs. the base system. Other applications of RNNs in medical texts have explored their performance in tasks such as deidentification of clinical notes^[33]^, information extraction^[44, 45]^, named entity recognition^[46]^, relation extraction^[47]^ and text classification^[48]^ with superior results to the reference models. However, we find did not find any articles that assessed the capacity of these models to classify clinical notes in Spanish.

### Aim

The main objective of our research is the comparison of two strategies for the classification of clinical notes in Spanish according to the patient’s smoking status: 1. a traditional approach involving task specific knowledge and heavy manual feature engineering and, 2. An ‘End-to-End’ approach. For the first strategy we used a bag of words representation, developed a task specific parser and trained two classifiers: a multi-layer perceptron (MLP) and a support vector machine (SVM). For the End-to-End strategy we implemented a deep bidirectional RNN using word embeddings as input.

### Dataset

Clinical notes were obtained from the EHR database at Hospital Italiano de Buenos Aires, Buenos Aires, Argentina. After taking a random sample of 6000 notes out of all the clinical notes from the ambulatory, emergency room and inpatient setting, we established that only 3.65% (95% CI 3.14 - 4.1) contained some information related to smoking. In order to optimize the manual labeling process given the low proportion of notes with information on tobacco use, we defined a filter based on four stems (‘tab’, ‘cig’, ‘fum’, ‘tbq’) to achieve a sample that had a greater proportion of notes with information on tobacco. We evaluated this filter on another set of 6000 notes sampled at random (which were also manually labeled), resulting in a negative predictive value of 0.99948 (95% CI 0.9985 - 0.9999) and a positive predictive value of 0.83077 (95% CI 0.7796 - 0.8743). Using this filter, we selected a sample of notes that contained any of the four stems from a random sample of 3054 patients 18 years of age or older, between 2005 and 2015. This yielded a total of 14718 clinical notes. The notes were labeled by three domain experts (family doctors) as ‘current smoker’, ‘current non-smoker’ and ‘clinical note without information on tobacco’ (inter-annotator agreement: kappa 0.97, 95% CI 0.9 - 1). The label ‘previous smoker’ was avoided since we found that in many cases these ex-smokers were described as non-smokers in the notes and that patients were almost never described as ‘never smokers’. So, the ‘current non-smoker’ category includes ‘never smokers’ and ‘ex-smokers’. Table 1 shows the characteristics of the dataset.

**Table 1.**
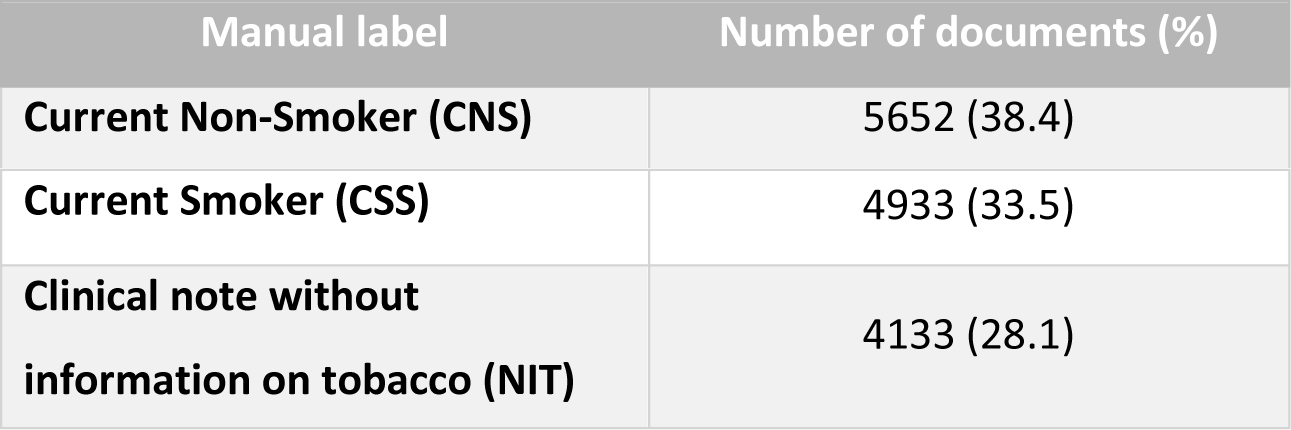
Dataset characteristics.

## Methods

Once the filtered sample of 14718 clinical notes was obtained, it was randomly split into a training set (11775 notes, 80%) and a test set (2943 notes, 20%). The test set was set aside and only used for the evaluation of the final models. The training set was used to train all three models, using 10-fold cross-validation for the hyperparameter tuning and feature selection processes. The final performance of the classifiers in the test set was assessed using per class and macro-averaged precision (positive predictive value), recall (sensitivity) and F1-score. All analyses were performed in R 3.4.1^[49]^ using the dplyr, ggplot, tidyr, text2vec, keras, tidytext, tensorflow and caret packages. All models were trained through the Keras API^[50]^ with TensorFlow^[51]^ as backend. Figure 3 shows an overview of the models’ development and validation process.

**Figure 3.**
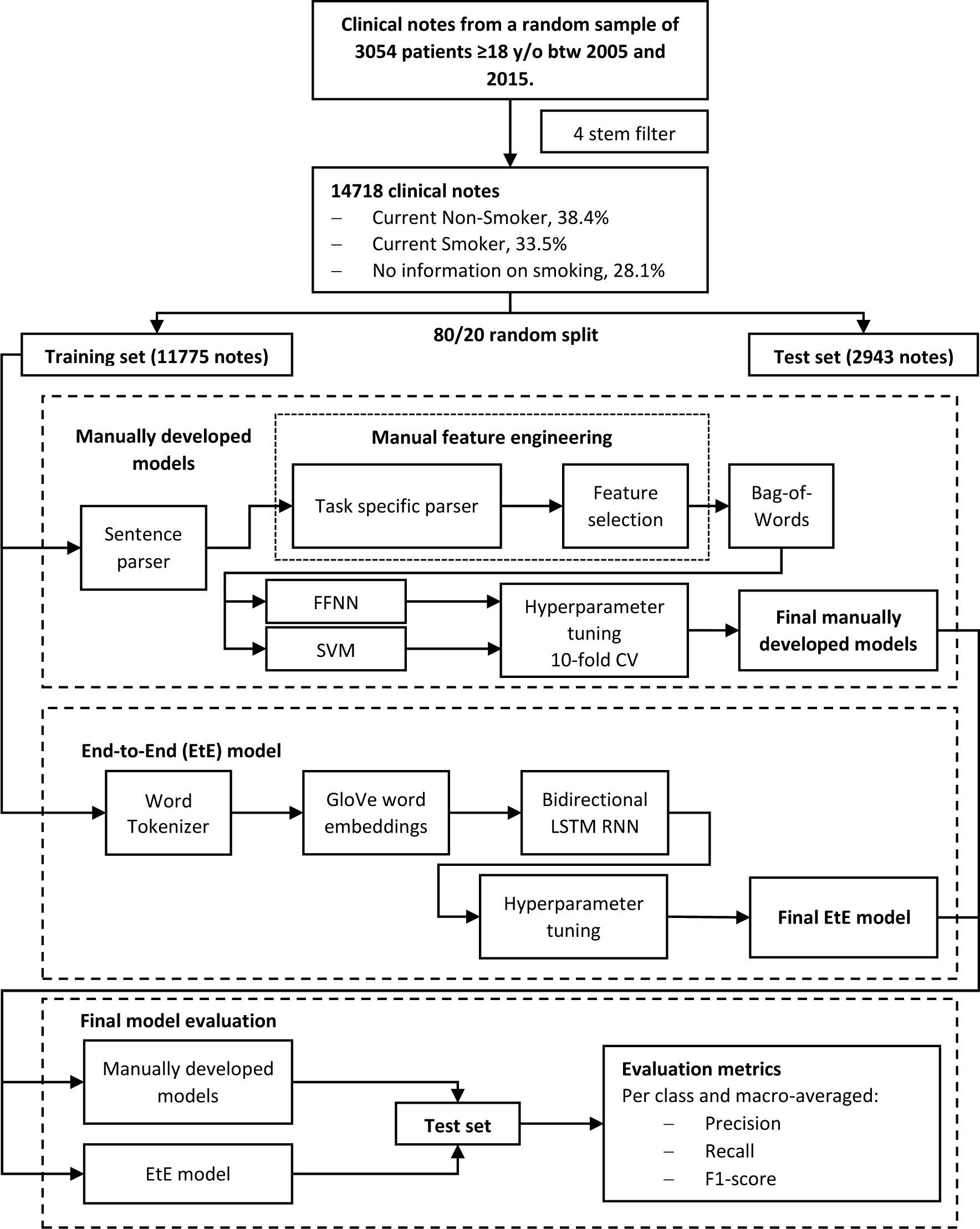
Overview of the models’ development and validation process. EtE: End-to-End model; MLP: Multi-layer perceptron; SVM: Support Vector Machine; CV: Cross-validation.

### Manually developed models

The development of the models was divided into two stages: 1. Development of a task specific parser and manual feature engineering, 2. Classifier training

#### 1. Task specific parser development (feature engineering and selection)

In order to incorporate task and context specific knowledge into the feature engineering process, we developed a task specific parser that allowed us to detect key smoking related terms, negations of those terms, positive and negative modifiers of those terms and modifier negation terms. Also, we detected who the subject associated with key smoking terms was.

Twelve key terms related to smoking were defined. The key term negation detection module was developed using regular expressions. This module detects negation terms 25 characters before and 10 after the key terms, unless some punctuation sign is detected such as commas, periods, question marks, semicolons and parentheses. Positive and negative key term modifiers were detected from a predefined list. Using a logic similar to that of the negation module, these modifiers were only detected if they were in the proximity of the key terms. Then the detection of negations of these modifiers was added. Finally, a list of terms that would detect other subjects related to the central terms (mother, father, brother, spouse, etc.) was developed. Figure 4 shows an example of the parser applied to a sentences containing information on smoking status.

**Figure 4.**
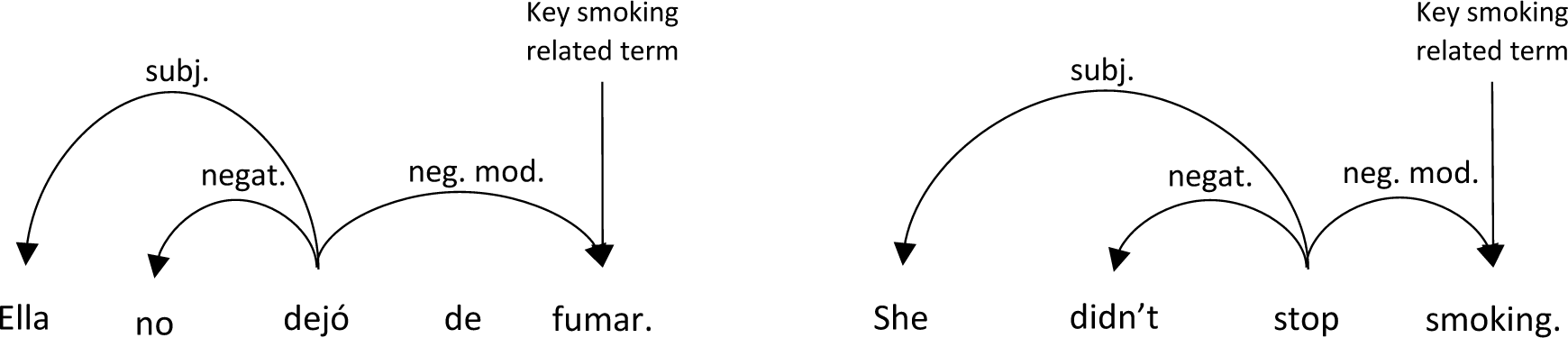
Example of the task specific parser (Spanish and English) *Subj:* subject; *negat.:* negation; *neg.mod.:* negative modifier

Once this parser was defined, the documents were segmented into sentences and only those that had any of the key smoking related terms were kept. A table was constructed in which each row was a sentence and each column contained the counts of each of the terms detected by the parser per sentence. Thus, initially, 9021 features were defined. This number was later reduced to 561 after dropping those with a spearman correlation coefficient greater than 0.85 and those that had an entropy-based importance of zero. Completing this process required 280 hours.

#### 2. Classifier training

For both classifiers, hyperparameters were evaluated using a grid search approach and 10-fold cross validation.

##### Multi-layer perceptron

The final classifier resulted in a multi-layer perceptron (feed-forward fully connected artificial neural network) with three hidden dense layers (483, 377 and 358 hidden units respectively) and a Softmax layer as the output layer (figure 5). It was optimized for 150 epochs in batches of 128 observations by mini-batch gradient descent with momentum of 0.9, learning rate of 0.2 and learning rate decay of 0.004. We used Rectified Linear Units (ReLU) as activation functions between layers and categorical cross entropy as the objective function. All hyperparameters were tuned using 10-fold cross-validation. L2 regularization and dropout^[52]^ were evaluated to avoid overfitting, but they did not have a significant impact on the model’s performance in the cross-validation set so they were not incorporated into the final model. The total training time of the classifier was 40 hours.

**Figure 5.**
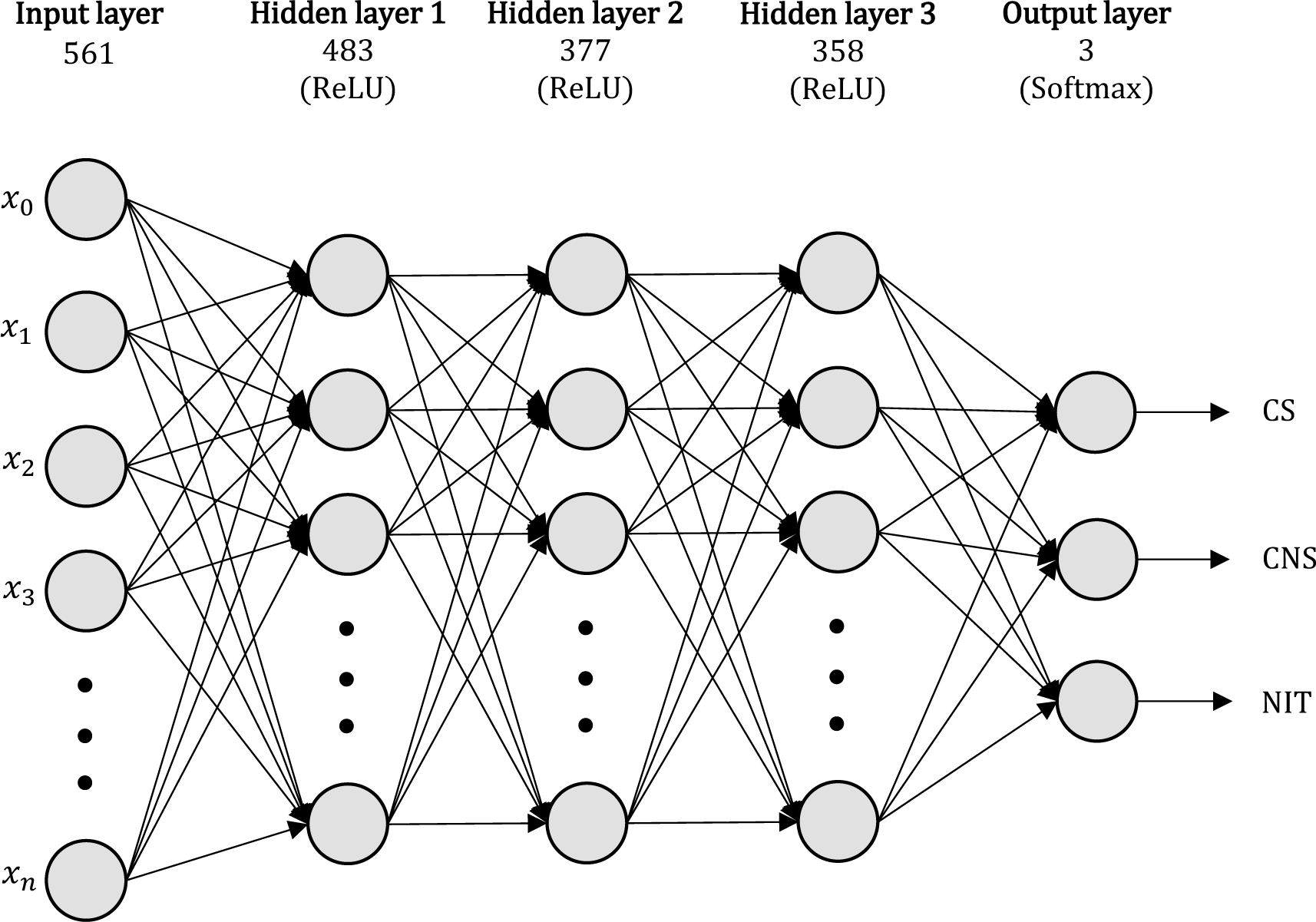
Multi-layer perceptron (feed forward neural network) CS: Current smoker; CNS: Current non-smoker; NIT: no information on tobacco; ReLU: Rectified linear unit; x: input.

##### Support vector machine

As an alternative to the MLP we trained a support vector machine^[53]^ on the same 561 features dataset as the MLP. We tested three different kernels: lineal, polynomial and radial. Finally, the radial basis function was selected with hyperparameters cost=8 and gamma=0.57. Total training time was 16 hours.

#### End-to-End model

##### 1. Classifier training

We trained a RNN at the document level with a word embedding layer, three hidden bidirectional LSTM layers (100, 100, 100 hidden units per layer per direction), three hidden dense layers (100, 100, 50 hidden units) and a Softmax layer to output the probabilities for each category (fig. 6). We used recurrent dropout^[54]^ (dropout probability = 0.2) to avoid overfitting in the LSTM layers and regular dropout (dropout probability = 0.2) in the dense layers. The activation function between the LSTM layers is tanh and ReLU between the dense layers. As optimization algorithm we used RMSprop^[55]^. We also used categorical cross entropy as the objective function. All hyperparameters were tuned using 10-fold cross validation. The total time of development of the model was 56 hours. The first 10 hours were used for the training and exploration of the word embeddings; the last 46 were assigned to training the network.

**Figure 6.**
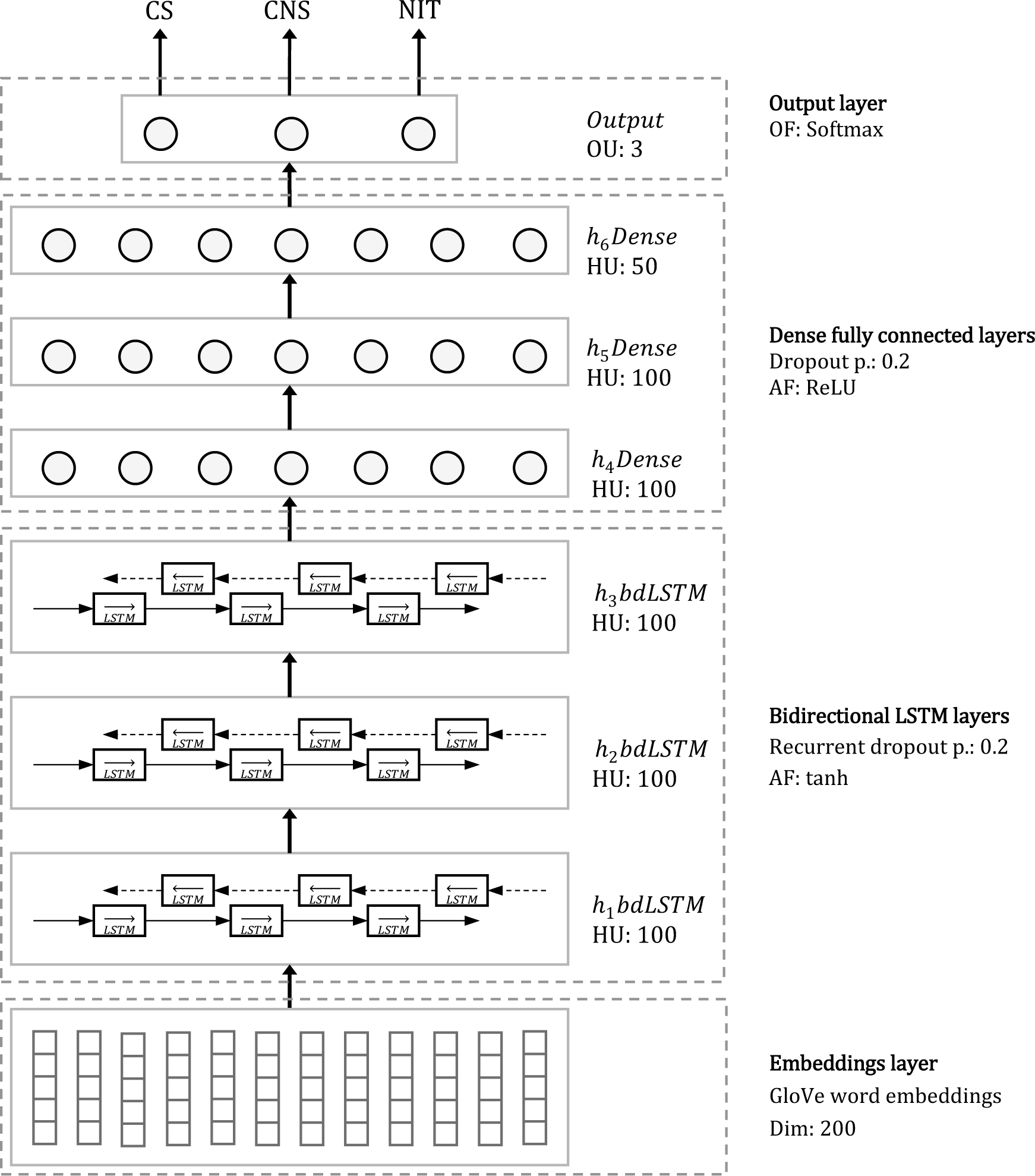
Final End-to-End model. *h*: hidden layer; *bdLSTM*: bidirectional long-short-term-memory layer; *HU*: Hidden units; *Dim*: dimensions; *Dense*: dense fully connected layer; *p*: probability of retention; *AF*: activation function; *ReLU*: rectified linear units; *OU*: output units; *OF*: output function, CS: Current smoker; CNS: Current non-smoker; NIT: No information on tobacco.

##### GloVe Word embeddings

In order to initialize the embedding layer of the RNN, we pretrained word embeddings from a corpus formed by the training set documents plus the 12,000 documents of the samples described in the *‘Dataset’* section, using the model described by Pennington et. al^[37]^. A window of 10 words was selected. Of the 36,558 unique words, those that appeared less than 5 times throughout the corpus were discarded. The remaining 9,483 tokens were trained to obtain 200 dimensional vectors.

## Results

Table 2 and figure 7a show the performance of the three classifiers on the test set. The RNN was slightly superior to the MLP in all three macro-averaged metrics (difference in recall: 0.009 [95%CI: -0.0007, 0.017], precision: 0.007 [95% CI: -0.0015, 0.019] and F1-score: 0.009 [95% CI: 0.0018, 0.016]), although the 95% confidence interval only excludes zero for the difference in F1 score (figure 7b). When comparing the performance of the bdRNN (bidirectional RNN) to the SVM, the SVM performs better in general (difference in recall: -0.007 [95%CI: -0.016, 0.0018], precision: -0.009 [95% CI: -0.018, 0.00015] and F1-score: -0.008 [95% CI: -0.014, -0.0017]), where also the F1 score difference confidence interval excludes zero.

**Figure 7a.**
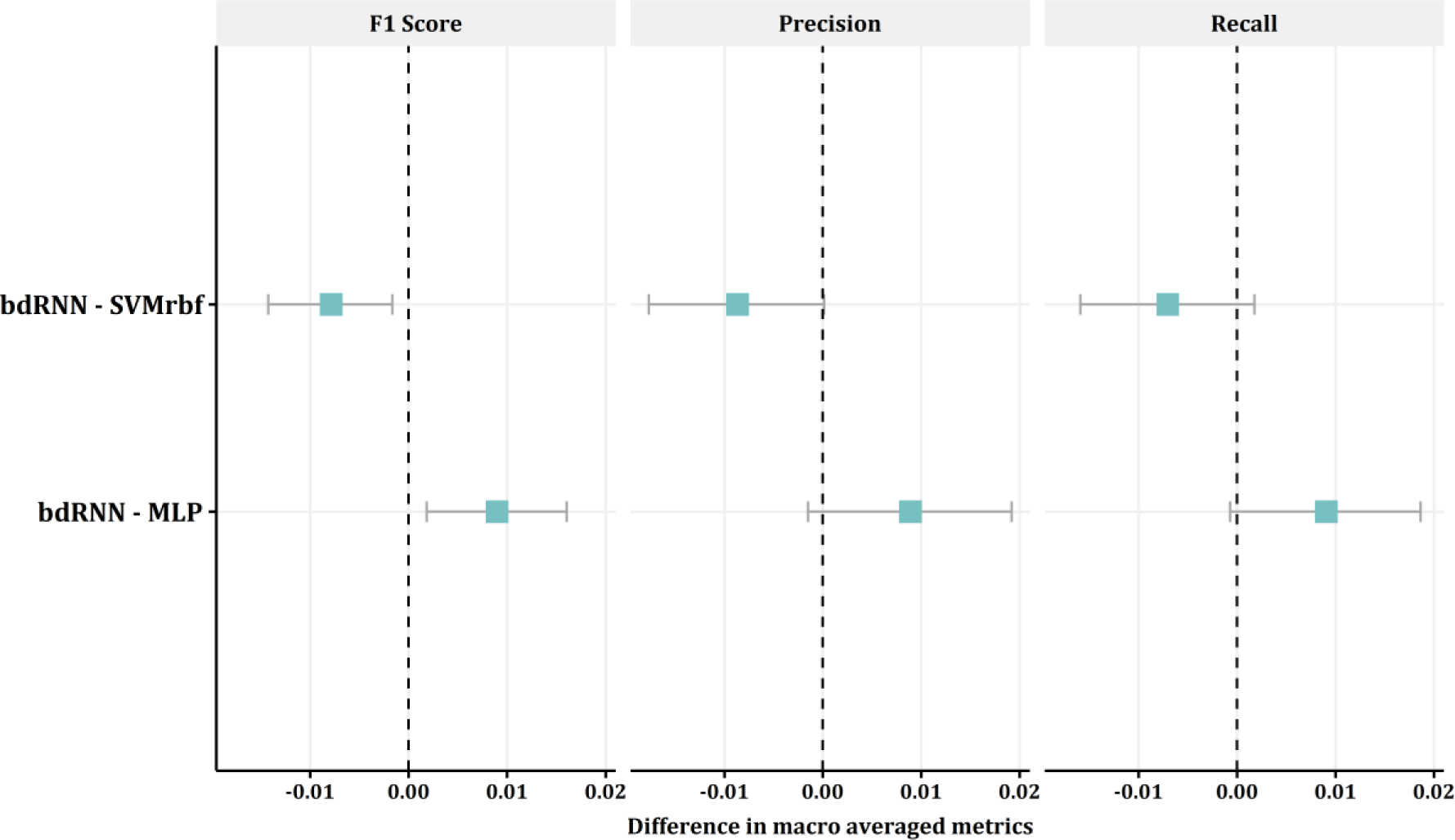
Performance of the classifiers on the test set.

**Figure 7b.**
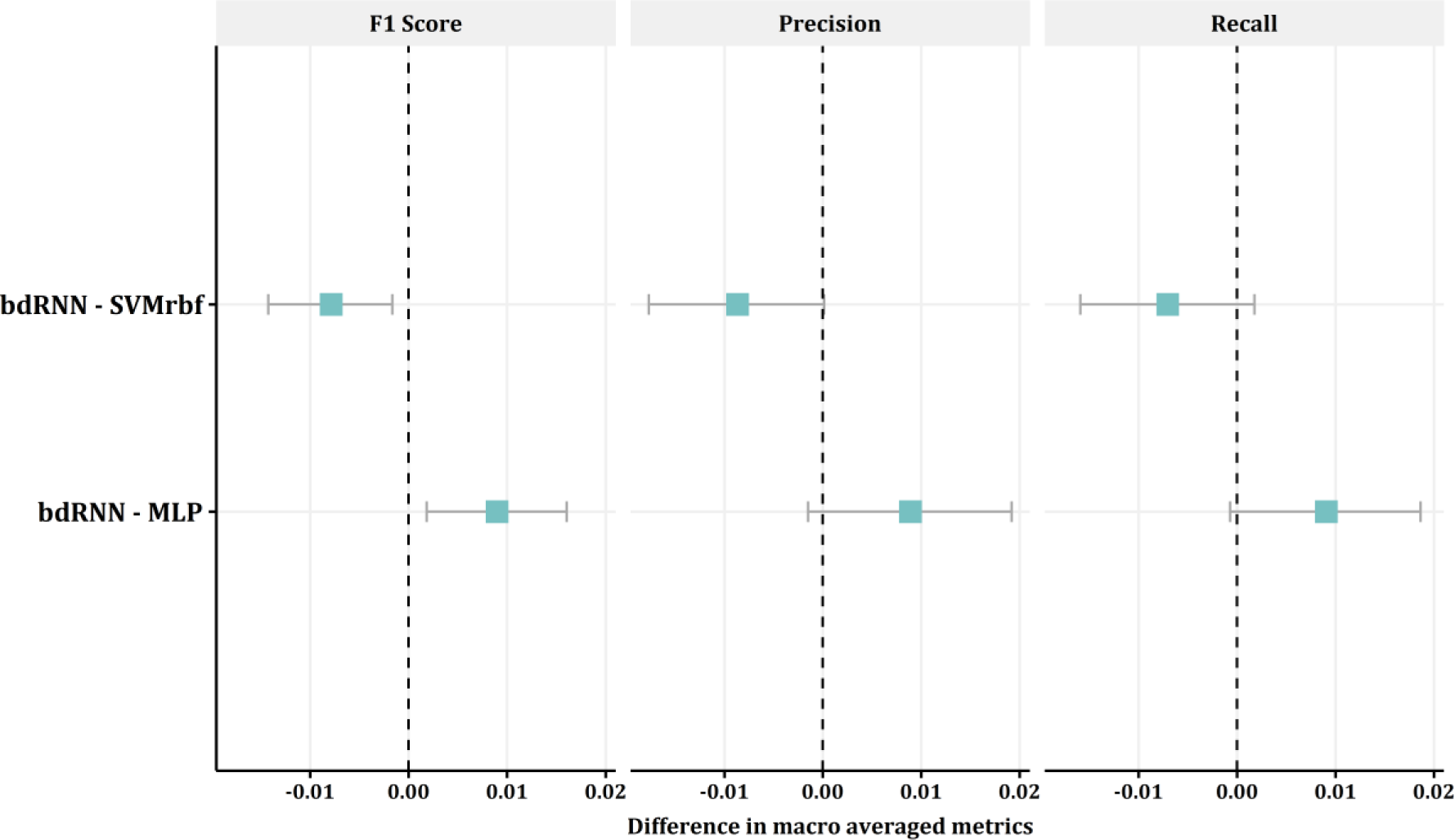
Difference in macro-averaged metrics between classifiers. Macro-averaged F1-score, precision and recall with 95% confidence intervals. bdRNN: bidirectional recurrent neural network; SVMrbf: support vector machine w/radial basis function; MLP: multi-layer perceptron.

**Table 2.**
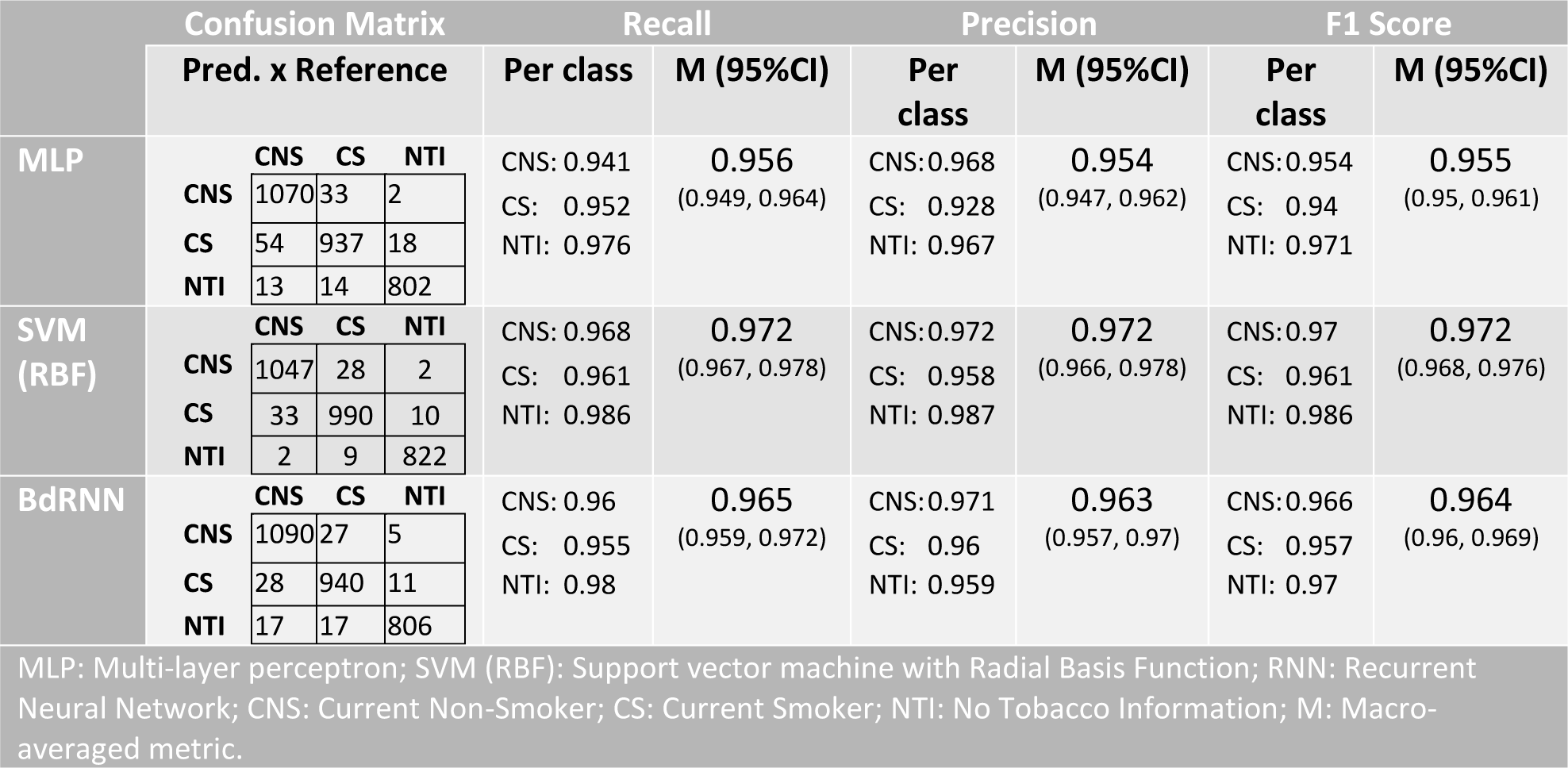
Performance of all three classifiers on the test set.

## Discussion

We developed a set of classifiers that detect smoking status from clinical notes is Spanish using a bag-of-word model with heavy manual feature engineering and a sequential End-to-End model by means of a bidirectional deep LSTM RNN. The End-to-End model performed similarly to methods that involve a great investment of time and task specific knowledge for the manual development of features. To the best of our knowledge, this is the first paper that evaluates the performance of these models in clinical notes in Spanish.

This capability of the RNN to represent the information contained in the documents without any manual development of features is explained by several of its characteristics. Bidirectional sequential processing allows the incorporation of previous and subsequent information to each word and is very similar to the way in which humans understand texts. On the other hand, the different layers of the RNN allow a hierarchical representation of the information, learning more abstract representations of the texts in each layer. This, added to the representation of the words by word embeddings that capture the semantic relationships between words, places this type of models as an excellent choice of End-to-End models.

Not having to engineer features manually facilitates texts analysis in languages or contexts for which NLP tools are not available. In addition, according to our experience, it saves time (in our case, the development of the RNN required less than 20% of the time it took to develop the manual models) and allows the analyst to focus on the architecture of the network rather than on the exploration of the notes and extraction of variables. This in turn makes the workflow easily exportable to other classification problems, in various contexts and languages unlike the process of manual variable development. In the particular context of public health management based on EHR data, it is even more valuable since it allows to potentially design, in a short time, precise classifiers for creating electronic phenotypes than can be used in policy evaluation or research. This, nevertheless, comes at the price of spending time in hyperparameter tuning and designing the structure of the network, which is usually a trial and error process.

Our work has several limitations. Among them, we use a set of bag-of-words models as baseline models since it is part of our usual workflow, however other alternatives such as conditional random fields or other state-of-the-art models could be excellent baseline models as well, as proposed by other authors^[33, 42, 44, 45, 47, 48]^. Also, we only evaluated the models in a classification problem with only a few classification categories. However, other teams have had very good results even with more categories and fewer observations^[42, 44]^.

## Conclusion

We have shown that deep bidirectional LSTM RNNs as End-to-End models achieve similar same levels of performance in the classification of clinical texts in Spanish, as models with heavy manual feature engineering albeit in less than 20% of the development time. This makes them an important tool for streamlining text processing in languages in which the development of NLP tools has not advanced as much as in English. Areas like research or public health management could clearly benefit from ‘End-to-End’ models that facilitate the exploitation of already available data sources such as EHRs.

